# Susceptibility of *Klebsiella pneumoniae* Clinical Isolates in Biofilms to Antibiotics and Assessment of Secondary Drug Effects

**DOI:** 10.64898/2026.05.15.725361

**Authors:** Daria A. Burmistrova, Nadezhda A. Gultiaeva, Kseniya V. Danilova, Ivan N. Kravtsov, Andrey I. Solovyev, Anastasia V. Kartashova, Olga L. Voronina, Marina S. Kunda, Natalia N. Ryzhova, Ekaterina I. Ermolova, Polyna V. Mazorchuk, Kristina A. Ryzhova, Lyubov’ A. Davydova, Vera Y. Baturova, Alexey I. Gutnikov, Irina V. Kolesnikova, Oksana V. Shelkovnikova, Yulia M. Romanova, Sergey V. Tsarenko, Alexander L. Gintsburg, Denis Y. Logunov

**Affiliations:** Department of Genetics and Molecular Biology of Bacteria, N.F. Gamaleya NRCEM, Moscow, Russia; Department of Bacterial Infection, N.F. Gamaleya NRCEM, Moscow, Russia; Department of Medical Microbiology, N.F. Gamaleya NRCEM, Moscow, Russia; Department of Infectology and Virology, I.M. Sechenov First MSMU, Moscow, Russia; Department of Clinical Pharmacology, National Medical Research Center, Center for Treatment and Rehabilitation, Moscow, Russia; Hospital Administration, National Medical Research Center, Center for Treatment and Rehabilitation, Moscow, Russia; Laboratory Department, National Medical Research Center, Center for Treatment and Rehabilitation, Moscow, Russia; ICU Department No. 3, National Medical Research Center, Center for Treatment and Rehabilitation, Moscow, Russia; ICU Department No. 2, National Medical Research Center, Center for Treatment and Rehabilitation, Moscow, Russia; Diagnostic Department, National Medical Research Center, Center for Treatment and Rehabilitation, Moscow, Russia; Faculty of Fundamental Medicine, Lomonosov Moscow State University, Moscow, Russia

## Abstract

Biofilms pose a significant challenge to antimicrobial therapy. Bacteria in biofilms differ from planktonic counterpart in their altered metabolism, collective behavior, protective role of extracellular matrix and diversified microbial subpopulations. These attributions significantly influence bioavailability and activity of antibiotics. The presence of bacterial aggregates during acute infections expands the problem to many other conditions previously not discussed in the biofilm context. *Klebsiella pneumoniae* is a leading cause of life-threatening hospital-acquired infections and is included in the WHO Bacterial Priority Pathogens List due to increasing antimicrobial resistance. The combination of antimicrobial resistance and the ability to form biofilms severely limits the efficacy of antibiotic treatments. In this study, we investigated the *in vitro* susceptibility of mature biofilms to 13 antimicrobials of *K. pneumoniae* clinical isolates from a single hospital. The resistance profiles of the local clinical isolates were consistent with the global epidemiology of *K. pneumoniae*. Minimal biofilm eradication concentrations (MBEC) for mature biofilms were defined with two assays (biomass and metabolic activity measurements) and brought into relation with susceptibility breakpoints and plasma (*C*_max_). Colistin sulfate, tigecycline, cephalosporins and combination of imipenem with cilastatin were the most potent biomass eradicators, while suppression of metabolic activity was barely reachable. Moreover, we observed a notable increase in metabolic activity upon exposure to sub-MBEC concentrations of antibiotics. Finally, our data broach a subject of antibiotic prioritization with respect to biofilm tolerance.

**IMPORTANCE:** This study addresses the critical gap between standard antibiotic susceptibility testing and the tolerance of biofilm and microbial aggregates during infections caused by *K. pneumoniae.* By systematically evaluating mature biofilms from a significant number of clinical isolates, we demonstrate that colistin and tigecycline show potent activity against both biofilm biomass and metabolic activity, whereas cephalosporins primarily reduce biomass without effectively suppressing bacterial metabolism, and other drugs have only weak effects on biofilms at clinically achievable concentrations. Furthermore, the alarming observation that sub-inhibitory biofilm eradication concentration (sub-MBEC) of antibiotic can paradoxically increase the metabolic activity of biofilms highlights a potential risk factor for therapy failure and resistance development. Our findings contribute to the necessary evidence base for prioritizing existing antibiotics in the limited armamentarium against biofilm-forming *K. pneumoniae*.

## INTRODUCTION

A biofilm is defined as a structured consortium of microbial cells surrounded by a self-produced polymer matrix, although some terminological uncertainty persists [1]. Depending on the system or environment, spatial and temporal variability within biofilms exists. Diverse conditions give rise to a wide range of structures – from simple microbial clusters and aggregates to highly organized and sustainable microbialites, as well as unusual forms such as DNA damage-induced R-structures in *Klebsiella pneumoniae* – all of which are considered biofilms [2, 3].

From a clinical perspective, bacterial biofilms tolerate antimicrobials and have a great impact on therapy failure despite adequate prescription according to standard laboratory testing. Biofilm tolerance is regarded as the most crucial characteristic in the context of infections; consequently, tolerance to antimicrobials and therapy resistance have been proposed as criteria to distinguish the biofilm lifestyle from planktonic clumps. Mostly biofilm infections are attributed to foreign body infections (i.e., ventilator associated pneumonia (VAP), catheter related infections) and chronic infections (i.e., cystic fibrosis). However, Kolpen et al. recently have demonstrated a presence of biofilms (microbial aggregates) during acute pneumonia (non-VAP) as well. According to this study the main difference in aggregates during chronic and acute lung infection is their metabolic activity [4]. The presence of bacterial aggregates during acute infections expands the problem to many other conditions previously not discussed in this context. Bacteria in biofilms differ from planktonic counterpart in their altered metabolism, collective behavior, protective role of extracellular matrix and diversified microbial subpopulations. These attributions significantly influence bioavailability and activity of antibiotics [5, 6].For example, biofilm extracellular matrix not only creates a physical barrier around bacterial community but also contains drug conversing enzymes and extracellular molecules acting as distracting targets for antibiotics on its path to bacterial cells [7, 8, 9, 10]. Within biofilm some bacteria persist in metabolically inactive state and survive prolonged exposure to antimicrobials [11]. Other subpopulations might die under the influence of antibiotics in that way increasing extracellular biofilm content and protecting deeper laying survivors. Notably, bacterial death upon exposure to antimicrobials can be either a passive process or a regulated ’kamikaze’ behavior [12]. Moreover, mutation rate, horizontal gene transfer (HGT) and development of resistance are intensified in biofilm state thus promoting multidrug-resistant (MDR) pathogen emergence [13]. Standard antimicrobial testing relies on planktonic culture of microorganisms while biofilms are outside the scope of clinical diagnostics [14] [1]. Several previous studies were devoted to minimal biofilm eradication/inhibition concentrations (MBEC / MBIC) of some common antibiotics.

Unsurprisingly, the reported MBEC values were substantially higher (often by orders of magnitude) than the corresponding minimal inhibitory concentrations (MIC) for clinical isolates of *Pseudomonas aeruginosa*, *Acinetobacter baumannii*, *Klebsiella pneumoniae*, *Escherichia coli*, *Staphylococcus aureus* [15, 16, 17, 18]. Despite well recognized role of *K. pneumoniae* as one of the leading causes of hospital acquired infections and member of ESKAPEE group of pathogens there is not much data on activity of common antibiotics against its biofilms. Comprehensive studies devoted to screening of *Klebsiella* biofilm susceptibility to antibiotics are limited and listed in Table **1**. Other relevant investigations are based on a very limited number of isolates [19, 20, 21].

**TABLE 1.** Characteristics of *K. pneumoniae* isolates and MBEC determination methodology across different studies.

| *K. pneumoniae* characteristics and source | Antibiotics | Methodology / Assay used | Sample size | Reference |
| --- | --- | --- | --- | --- |
| KPC-producing XDR clinical isolates isolated in the Tel-Aviv Sourasky Medical Center before 2014, Israel | gentamicin, colistin | Turbidity / regrowth assay | 46 | [26] |
| XDR and PDR colistin-resistant clinical isolates isolated in 2016-2019 at the King Chulalongkorn Memorial Hospital, Bangkok | colistin, colistin-EDTA combination | Crystal violet, PrestoBlue<sup>1</sup> | 47 | [27] |
| Clinical isolates isolated in 2017 at the Affiliated Hospital of Southwest Medical University in Sichuan province, China | cefepime, amikacin, ciprofloxacin, imipenem, polymyxin B | Turbidity / regrowth assay | 16 | [28] |
| Carbapenem-resistant strong biofilm-forming hypervirulent (hv) isolates isolated from bloodstream infections archived at the Department of Clinical Microbiology, Christian Medical College, Vellore, India during 2019-2020 | minocycline, colistin, ceftazidime/avibactam, ceftazidime/avibactam with aztreonam, cefepime-zidebactam | Not specified | 16 | [18] |
| Clinical isolates isolated in 2023 at the National Medical Research Center "Treatment and Rehabilitation Center", Moscow, Russia | amikacin, gentamicin, meropenem, ertapenem, imipenem/cilastatin, colistin sulphate, tigecycline, cefepime, ceftriaxone, ceftazidime/avibactam, levofloxacin, moxifloxacin, ciprofloxacin. | Crystal violet, Resazurin assay | 51 | This study |

XDR – extensively drug-resistant; PDR – pan drug-resistant; KPC – *Klebsiella pneumoniae* carbapenemase.
<sup>1</sup>PrestoBlue® assay is based on resazurin chemistry, the manufacturer does not disclose the exact concentrations or composition.

There are no specific antibiofilm drugs studied in clinical trials [22, 23]. So close-in perspective must rely on prioritization and repurposing of common antibiotics which are active against both planktonic and biofilm embedded bacteria [24, 25]. Here we present an *in vitro* study of antibiofilm activity of common antibiotics against mature biofilms of clinical isolates of *K. pneumoniae*, mostly with MDR phenotype. For measuring antibiofilm activity we have applied two methodologies – (1) total biomass staining with crystal violet and (2) measuring of biofilm metabolic activity with resazurin. Analysis of the data in conjunction with known pharmacokinetics parameters let us to speculate in potential prioritization of some drugs in the case of infection due to biofilm forming *K. pneumoniae*.

## MATERIALS AND METHODS

### Klebsiella pneumoniae isolates

52 isolates were obtained during 2023 from a single hospital - National Medical Research Treatment and Rehabilitation Center of the Ministry of Health of Russia (Moscow, Russia). Isolation and initial species identification were made as a part of clinical diagnostic procedure using standard microbiological culture techniques. Isolates were deposited in laboratory collection as suspension in TSB medium supplemented with 15% glycerol and stored at -80°C.

### Matrix-assisted laser-desorption/ionization time-of-flight (MALDI-TOF)

Bacterial culture was seeded from glycerol stock on TSA plate and incubated for 24-48 h at 37°C. At least 3 single colonies for each isolate were directly transferred on stainless plate, extracted with 1 mkl 70% formic acid and covered with 1 mkl HCCA matrix. After drying plate proceeded to Biotyper® analysis on Ultraflex MALDI (Matrix Assisted Laser Desorption/ Ionization) tandem mass spectrometer (Bruker, Germany) or ALMASS Bio 200 (Algimed Techno, Belarus). Only confident level of identification (score value >2.0) was accepted.

### Antibiotic susceptibility testing

Antibiotic susceptibility was done with disc diffusion method (except colistin testing) according to National standard and EUCAST 2024-01-01 guideline. For colistin susceptibility testing bacterial suspension was inoculated into Mueller-Hinton broth supplemented with colistin sulfate 2 mg/L and incubated for 18-24 h at 37°C. Breakpoints were defined as in v. 14.0 (valid from 2024-01-01) EUCAST.

### Biofilm formation

Overnight bacterial culture was diluted 1 to 100 in Mueller-Hinton broth and incubated at 37°C until mid-exponential growth phase, then inoculum was prepared with McFarland=0.5. Biofilm was grown in 96-well, clear, polystyrene, flat bottom, TC-Treated Microplates at 37°C, in humid atmosphere for desired time: 96 h for investigation of biofilm forming capacity of isolates; 48 h before addition of antibiotic in MBEC assay.

### Biofilm total biomass staining with crystal violet

After biofilm formation planktonic cells and conditioned medium were washed away; biofilm was fixed for 30 min with 96% ethanol followed by ethanol removal, overnight drying and staining with 0,1% solution of crystal violet in water for 30 min. Plates were washed by dipping them in a bath of tap water three times, when dried (typically for 24 h) and dye was extracted with 30% solution of glacial acetic acid. Optical density was measured at wavelength 596 nm (OD_596_) at BMG Clariostar® multi-mode microplate reader.

### Biofilm metabolic activity staining with resazurin

After biofilm formation conditioned medium were replaced with fresh medium supplemented with resazurin sodium salt at final concentration of 10 mkg/ml. Plates were incubated for 45 min at 37°C. Fluorescence of resarufin was measured with Ex/Em 540*±*20 nm/ 590*±*20 nm, top measurement with automatic gain adjustment at BMG Clariostar® multi-mode microplate reader.

### Minimal biofilm eradication concentration (MBEC) assay

Mature 48 h old biofilm were washed from planktonic cells and medium were replaced with antibiotic serial two-fold dilutions in fresh cation adjusted Mueller-Hinton broth. After the next 48 h of incubation plates were stained with crystal violet for biomass evaluation and with resazurin for bacterial metabolic activity (viability) assay. All points were performed in triplicate. MBEC was calculated as:

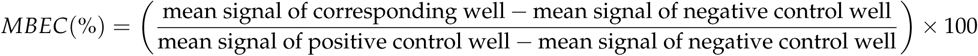

were positive control represents intact biofilm and negative control represents wells without biofilms. MBEC50 is a concentration causing at least 50% reduction in comparison with unexposed to antibiotic control. The choice of MBEC50 rather than MBEC75/90 was to reduce error due to asymptotic curves especially actual for CV staining. Assay agreement was calculated as percentile of isolates with reachable MBEC with both CV and resazurin from number of isolates with reachable MBEC with either CV or resazurin.

### Whole-genome sequencing and characterization

Whole-genome sequencing was performed on 15 *K. pneumoniae* isolates. The NadPrep EZ DNA Library Preparation protocol (Nanodigmbio (Nanjing) Biotechnology Co. Ltd., Nanjing, China) was used for the preparation of the library. Prepared libraries were sequenced with an Illumina NextSeq 550 sequencing platform. Genomes were assembled using CLC Genomics Workbench v. 21 (QIAGEN) and SPAdes v. 3.13.0 (St. Petersburg genome assembler, St. Petersburg, Russia, URL: ablab.github.io). The NCBI Prokaryotic Genome Annotation Pipeline (PGAP) was used for genome annotation. WGS data are available in GenBank: BioProject PRJNA561493, Accession Numbers are presented in supplementary Table 2. The spectrums of antimicrobial resistance genes were determined using the CARD (Comprehensive Antibiotic Resistance Database, URL: card.mcmaster.ca) resource and BV-BRC (Bacterial and Viral Bioinformatics Resource Center, URL: www.bv-brc.org). To clarify the annotation of beta-lactamases, the beta-lactamase database (BLDB, URL: bldb.eu) was consulted. Multilocus Sequence Typing (MLST) was performed with MLST tool (the BIGSdb-Pasteur platform, URL: bigsdb.pasteur.fr/klebsiella/).

## Results

### General description of studied clinical isolates of *K. pneumoniae*

All isolates were collected during 2023 year as a part of standard microbiological diagnostic procedure for patients with pneumonia (including ventilator associated pneumonia), urinary tract infections, catheter related infections, peritonitis, sepsis and others (supplementary table 1). Species identification was made on the basis of bacteriological culture technics and confirmed with MALDI-TOF MS. Antibiotic susceptibility for clinical purpose was defined with disc diffusion method, except colistin sulfate (Fig. **1**).

**FIG 1.**
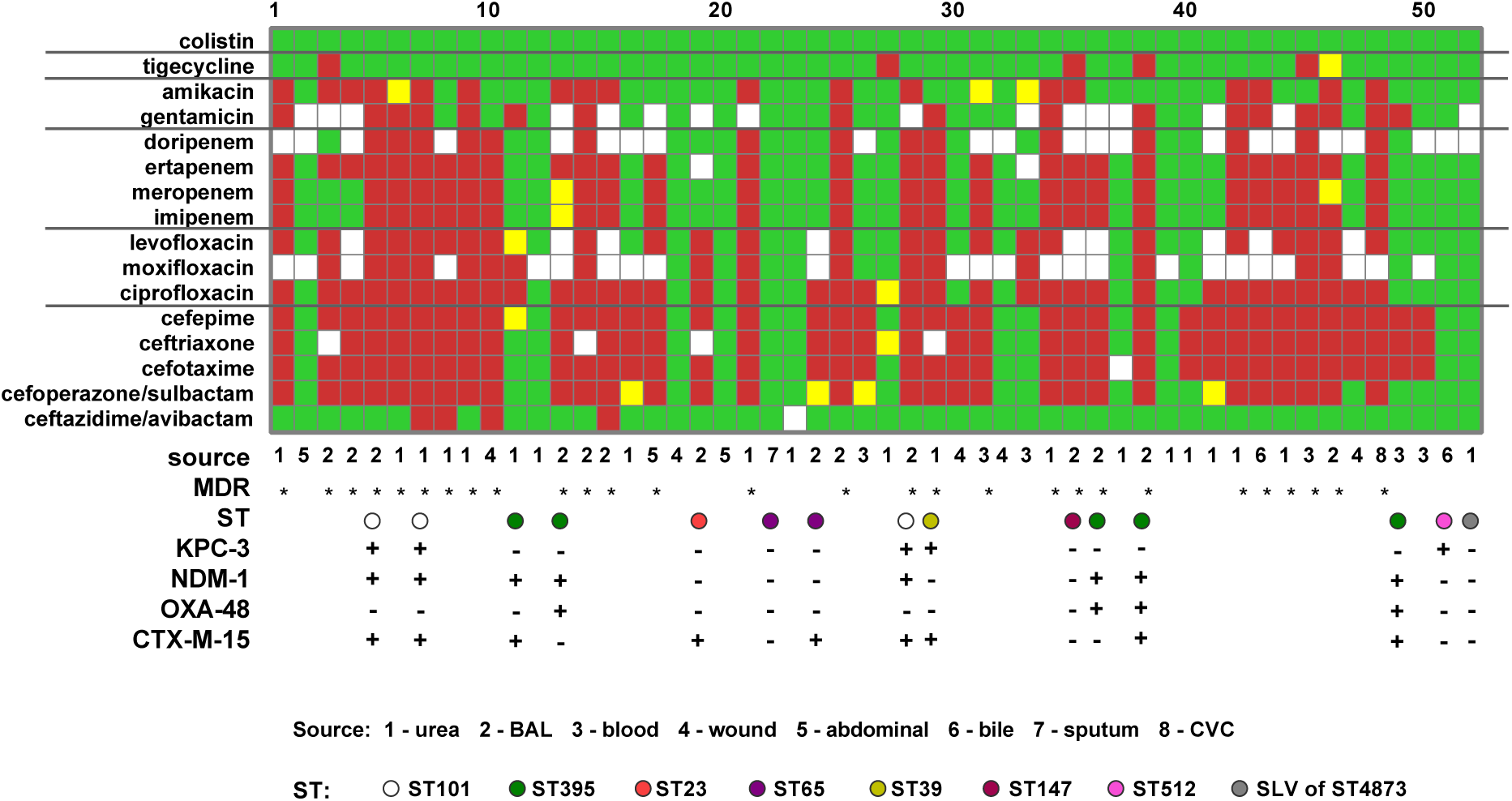
*K. pneumoniae* antibiotic susceptibility, sequence type diversity and presence of major carbapenemase and cephalosporinase genes. Green color - sensitive isolate, yellow - intermediate, red - resistant, white - not tested, * - multi drug resistant (MDR). Lower panel represents number-coded isolation source and color-coded sequence types (ST) according to the legend. Presence of KPC-3, NDM-1, OXA-48 and CTX-M-15-encoding gene is marked with a plus. Every tenth isolate is marked with a serial number at the top of the figure; the numbering order corresponds to the ranking by biofilm-forming ability, as in (Fig. 2). BAL - bronchoalveolar lavage; CVC - central venous catheter.

Isolates resistant to > 3 classes of antibiotics were classified as multidrug-resistant (MDR). In total 28 MDR *K. pneumoniae* were included in the study. Prevalence of carbapenem resistance (at least to one tested drug) was 53.8%, while tigecycline and ceftazidime/avibactam-resistant isolates were numerous and all isolates were sensitive to colistin.

### Clinical isolates of *K. pneumoniae* have diverse genetic background

The analyzed isolates were divided into eight genotypes ((Fig. **1** and supplementary table 2), seven of which are well known. *K. pneumoniae* ST147 belonged to the globally distributed clone, while *K. pneumoniae* ST101, ST23, ST39, ST395, and ST512 belonged to intercontinental clones. *K. pneumoniae* ST101, ST23, ST395, and ST512 were detected in Europe and Asia, while *K. pneumoniae* ST39 was revealed in Europe and Africa. *K. pneumoniae* ST65 was predominantly distributed in Asia. The genomes of ten isolates contained carbapenemase genes (KPC-3, NDM-1, and OXA-48) in various combinations. Seven isolates also contained CTX-M-15 cephalosporinase genes. The genomes of two isolates encoded only cephalosporinases. The genes of the listed beta-lactamases were absent in the genomes of three isolates.

### Clinical isolates of *K. pneumoniae* form biofilms *in vitro*

The ability to form biofilms *in vitro* has been studied for all clinical isolates. Static biofilms were grown for 96 hours in a liquid medium in 96 well plates, then washed from planktonic cells and stained with crystal violet (CV), since criteria for classifying bacteria by their ability to form biofilms were proposed for this test. The results ranked according to the obtained data are shown in (Fig. **2**).

**FIG 2.**
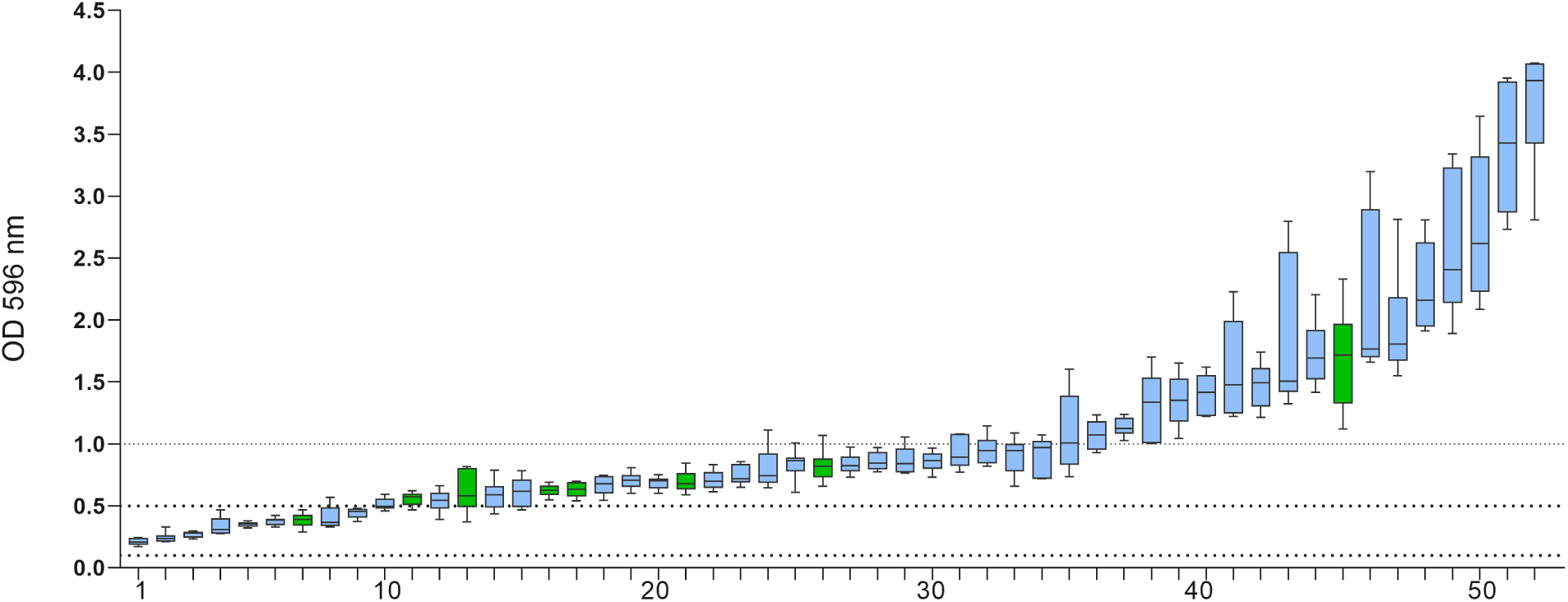
Ability to form biofilms *in vitro*. Biomass staining with crystal violet. Data represented as boxplots with median and whiskers from min to max. Green color represents hypermucoid isolates according to String test. Dotted lines set thresholds for weak, moderate and strong biofilm forming capacity classification. Every tenth isolate is marked with a serial number at the top of the figure; the numbering order corresponds to the ranking by biofilm-forming ability.

Overall, the majority of clinical isolates in our setting can be classified as moderate biofilm formers, whereas strong biofilm formers accounted for approximately 30%.

Although, as noted above, variability in biofilm formation may be observed depending on the conditions, we used the same methodological approach (plastic, medium, time, temperature) to assess the ability of clinical isolates to form biofilms as we used to evaluate tolerance to antimicrobial therapy.

### Biofilms display different level of tolerance to multiple classes of antibiotics

Minimal biofilm eradication concentration (MBEC) is a quantitative measure; therefore, it requires a defined threshold to determine the endpoint. In this study, we adopted the most commonly used criterion, MBEC50 (a 50% reduction compared to the untreated control), to assess biofilm susceptibility. In most studies researchers rely on 50% eradication (i. e. MBEC50). Other thresholds like MBEC75 or MBEC90 are not common partially due to level of measurement deviations, high background and non-linear resolution for CV and resazurin assays at low absolute units, so we also choose MBEC50 as a threshold to define biofilm susceptibility to antibiotics. Some data on antibiotic susceptibility were absent in clinical records (white square in Fig. **1**), so we experimentally restored missing data (supplementary table 1). Mature 48 h old biofilms were washed from planktonic cells and exposed to serial dilutions of antibiotics in a fresh medium for following 48 h, then stained with either CV or resazurin. In our experimental setup, the medium was replaced for 45 minutes for the resazurin assay, which is insufficient time to restore metabolism, regrowth and produce additional biomass. The number of isolates that responded to treatment in the biofilm state is presented in Table **2** and Figure **3**.

**FIG 3.**
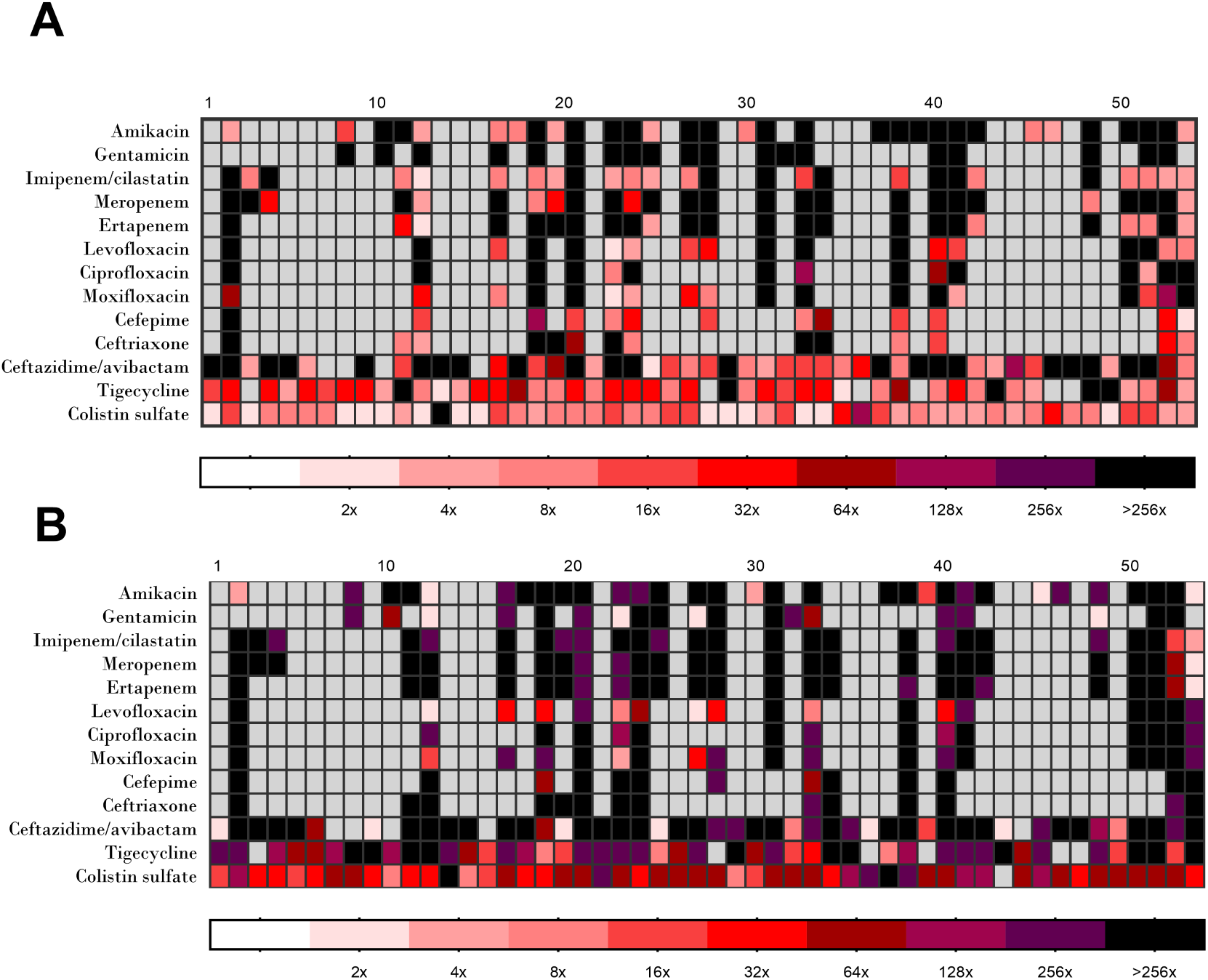
Minimal biofilm eradication concentration (MBEC50) against mature biofilms of antibiotic-sensitive *K. pneumoniae*. A – MBEC50 for biomass reduction, B – MBEC50 for metabolic activity suppression. Color intensity represents fold difference in MBEC50. Concentrations are expressed in folds over breakpoint for sensitivity of planktonic culture according to EUCAST (i.e., 1x/2x/4x/16x/32x/64x/128x/256x above breakpoint MIC). Black color represents cases where MBEC50 was not reached (more than 256x), grey color – resistant isolates, for which data on MBEC50 are hidden due to limited significance for clinical therapy. Every tenth isolate is marked with a serial number at the top of the figure; the numbering order corresponds to the ranking by biofilm-forming ability, as in Figure 2)

**TABLE 2.**
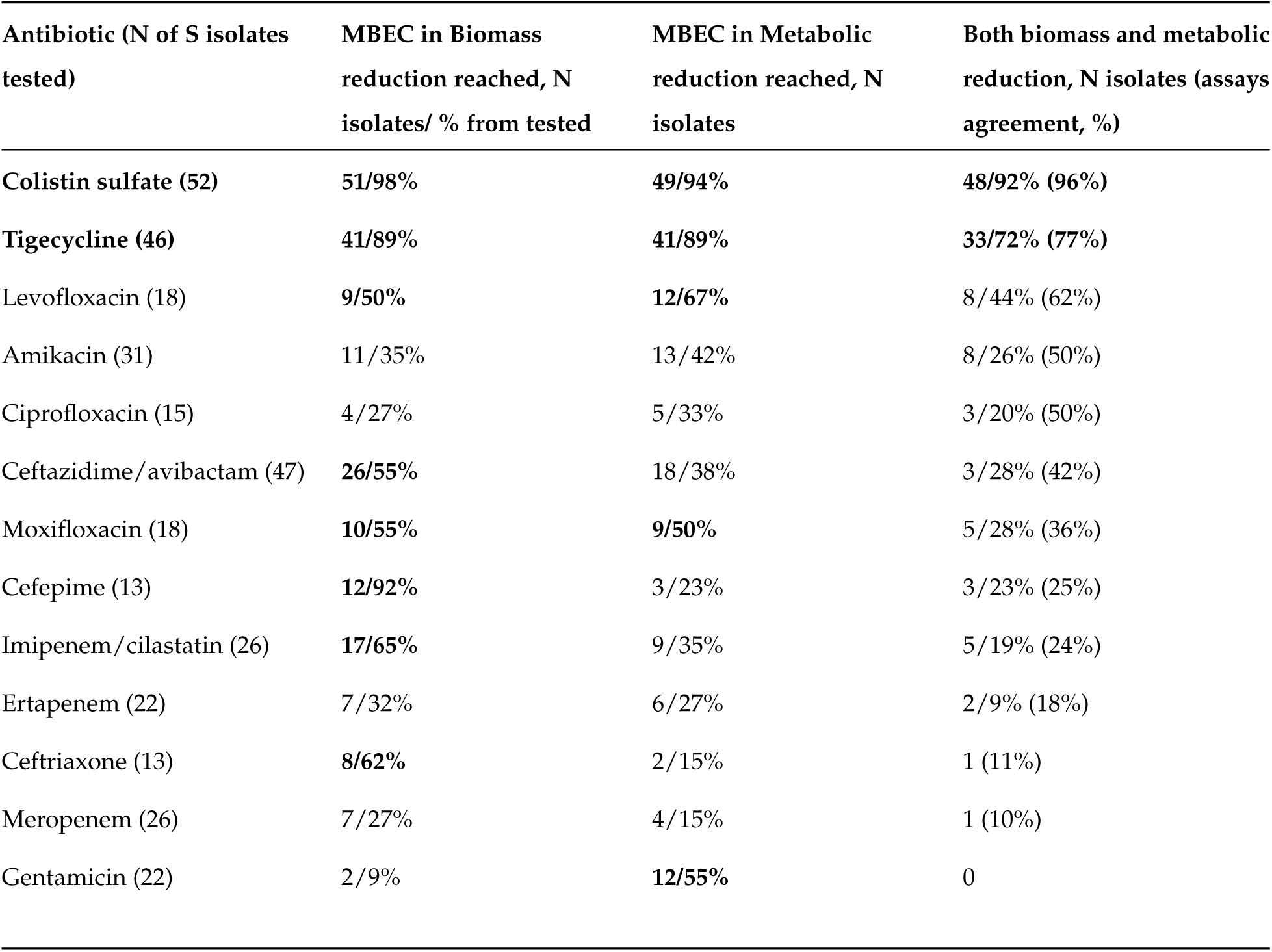
Biofilm responsiveness to different antibiotics. Data from two assays: CV staining of biomass and resazurin metabolic assay.

In general, there was a relatively low agreement between resazurin and CV assays.

Colistin sulfate and tigecycline caused reduction in both biomass and metabolism in exposed biofilms with high agreement between assays, but action of other drugs was multidirectional. Gentamicin caused reduction in metabolism, but had no effect on biomass. Cephalosporins showed trend in biomass reduction rather than metabolism suppression.

### Conjunction of antibiofilm potency with therapeutically reachable concentrations

The range of tested antibiotic concentrations was from 1x to 256x the breakpoint concentration for sensitivity definition (S according to EUCAST). While many studies compare MBEC and MIC, we propose comparing MBEC50 with the susceptibility breakpoint for sensitivity definition (S according to EUCAST) as a measure connected to clinically accepted criteria (see the lower right corner on Figure **4**).

**FIG 4.**
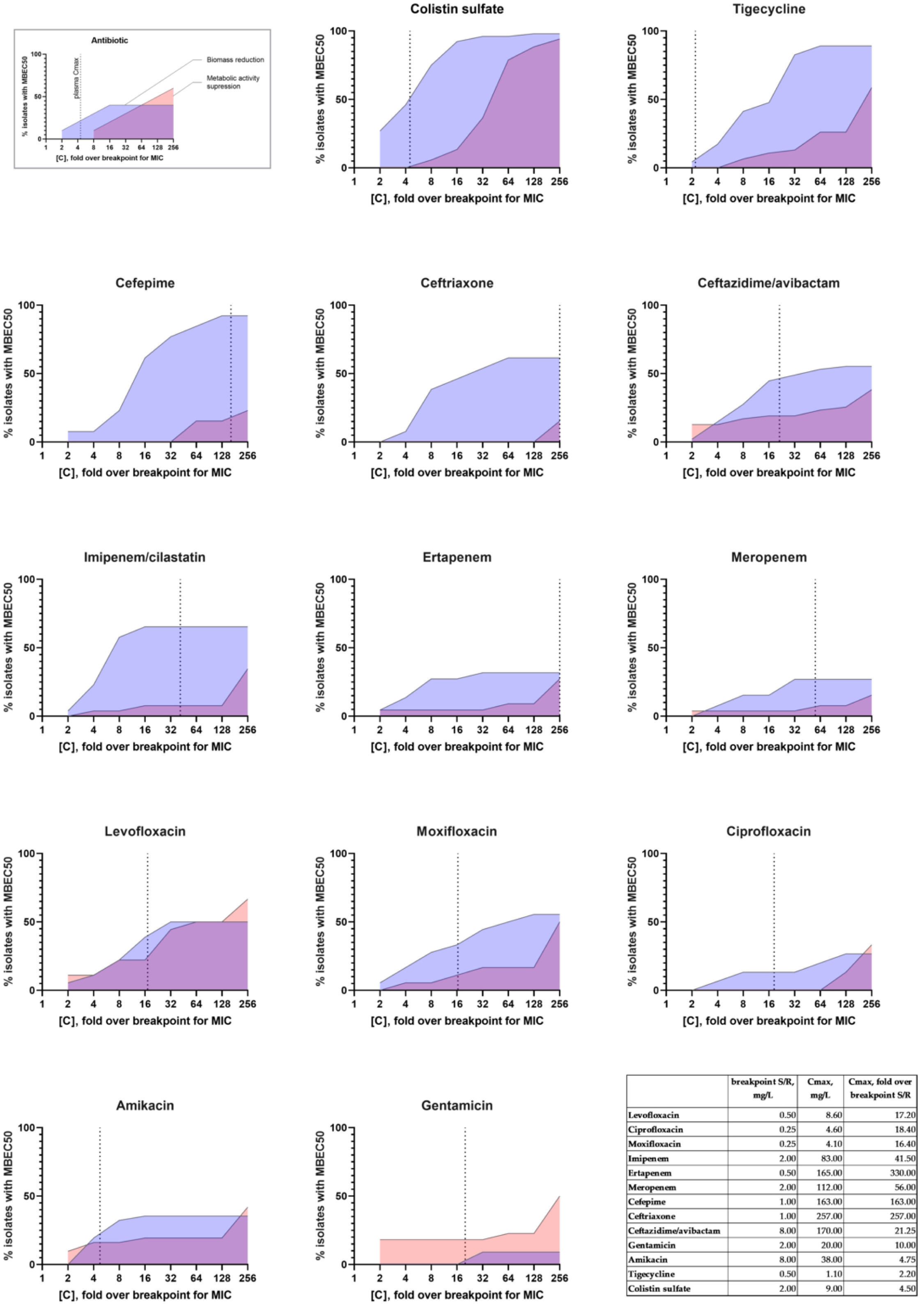
Successful rate in reaching MBEC50 in conjunction with plasma *C*_max_. The upper left corner contains a legend for reading the graphs. Vertical dotted lines represent plasma *C*_max_. X axis represents concentrations in folds against cutoff for sensitivity of planktonic culture according to EUCAST (i.e., 1x/2x/4x/16x/32x/64x/128x/256x above cutoff MIC). The lower right corner contains a table with *C*_max_.

While concentrations up to several orders above threshold for S/I/R MICs are usually tested in biofilm research it seems impossible to reach such concentrations during human therapy. The maximum plasma concentration (*C*_max_) is a widely accepted parameter to predict antibiotic efficacy (i.e., *C*_max_/*MIC* and other interconnected indexes) [29]. So for data analysis we additionally focused on plasma *C*_max_ reported in instructions for human use (intravenous rout of administration and the highest approved dose). To define if MBEC50 are reachable *in vivo* we have analyzed data in conjunction with plasma (*C*_max_). Chances to reach MBEC50 presented in figure **4**. It is worthy to note that for colistin sulfate, tigecycline and amikacin plasma *C*_max_ is not easily applicable in pharmacokinetic (PK)-based prediction of antibiotic efficacy. Most of tigecycline is distributed throughout the tissues after intravenous infusion, so plasma *C*_max_ is not representative for extrapolation. Colistin is a metabolite of prodrug sodium colistimetate, so pharmacokinetic/pharmacodynamic (PK/PD) is complex. Recommended target *C*_ss,avg_ = 2 mg/L is based on S/R cutoff rather than effective PK/PD indexes [30]. For amikacin *C*_max_/cutoff MIC (38/8 = 4.75) is lower than 8-10.

As «notable activity against biofilm» we have suggested at least 50 % chance (in terms of isolates number) to reach MBEC50. Plasma *C*_max_ for colistin and tigecycline is not useful for efficacy prediction, so we did not apply *C*_max_ in interpretation of MBEC for these antibiotics. With such assumptions colistin sulfate and tigecycline have the notable potencies in both biomass eradication and metabolic suppression. Cephalosporins and combination of imipenem with cilastatin have the notable potency in eradication of biofilm biomass at concentrations below plasma *C*_max_, but not in supression of metabolic suppression.

### Effect of subMBEC concentration on biofilm

While analyzing the metabolic suppression data, we observed an increase in metabolic activity for some isolates at sub-MBEC antibiotic concentrations. To ensure the robustness of our observations, an increase in metabolic activity was considered significant only if it was observed in at least two consecutive dilutions. Frequency of increased metabolism at subMBEC concentrations is presented in Table **3**.

**TABLE 3.** Increase in metabolic activity of biofilms under exposure to subMBEC of antibiotics.

| Antibiotic (N of S isolates tested) | Increase of metabolic activity at subMBEC,N isolates / % |
| --- | --- |
| **Colistin sulfate (52)** | **15/29%** |
| Tigecycline (46) | 6/13% |
| Levofloxacin (18) | 4/22% |
| Amikacin (31) | 4/13% |
| Ciprofloxacin (15) | 1/7% |
| Ceftazidime/avibactam (47) | 14/30% |
| Moxifloxacin (18) | 4/22% |
| Cefepime (13) | 4/30% |
| Imipenem/cilastatin (26) | 6/23% |
| Ertapenem (22) | 6/27% |
| Ceftriaxone (13) | 4/30% |
| Meropenem (26) | 5/19% |
| Gentamicin (22) | 6/27% |

Increase of metabolic activity was the most notable for colistin sulfate (partially due to the largest sample size), but other antibiotics also caused similar effect. As colistin sulfate was one of the most potent antibiofilm drug additional data presented in Figure **5**.

**FIG 5.**
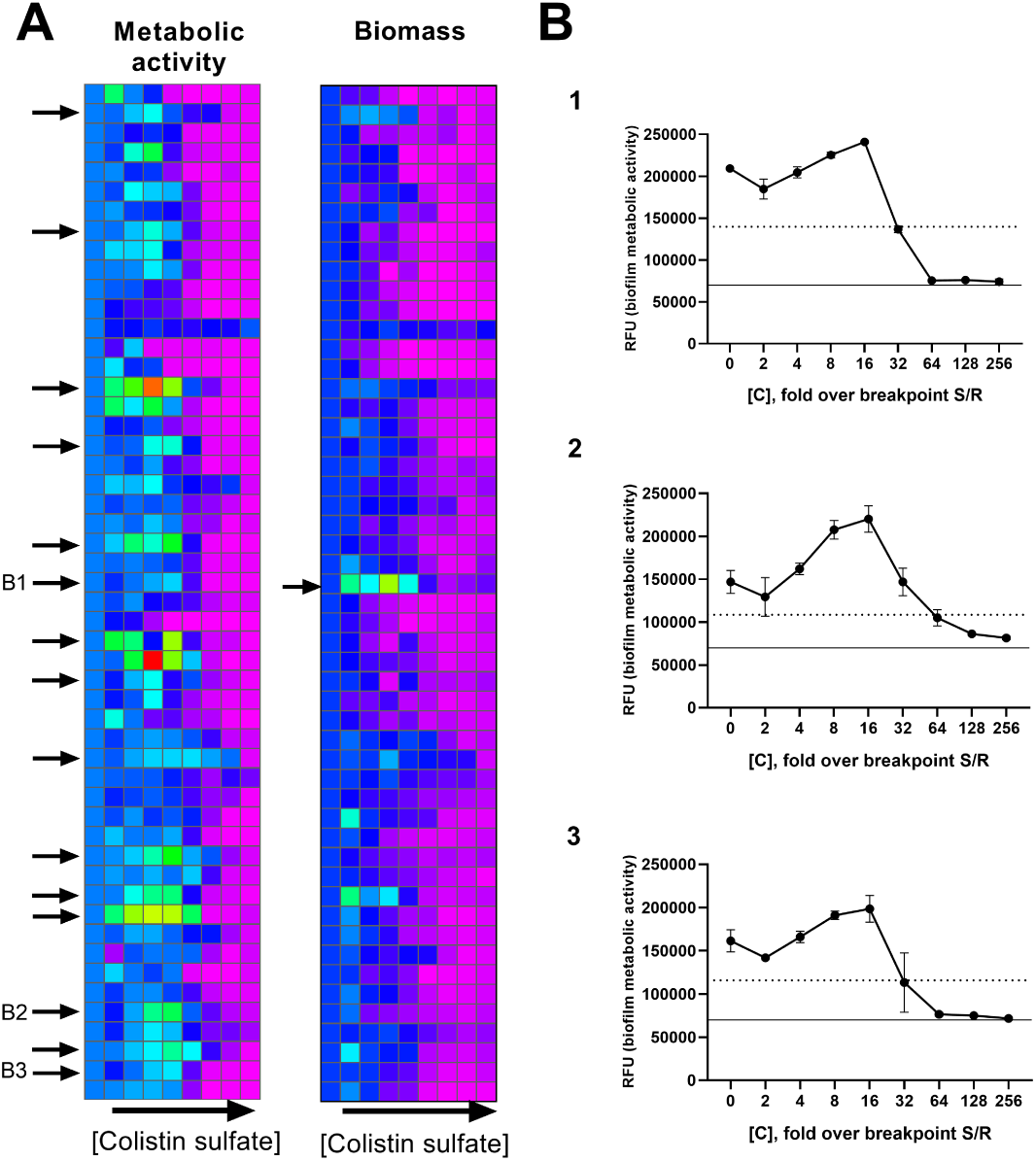
Increase in metabolic activity of biofilms under exposure to subMBEC of colistin sulfate. A – heat plot of normalized metabolic activity (left) and biomass (right) under exposure to colistin sulfate, arrows mark isolates with increase of metabolic activity/biomass above intact biofilm. B – representative examples of primary measurements of metabolic activity, dotted line – MBEC50, solid line – background control, mean and SD from 3 replicates.

Moreover, we exclude from our criteria effect found in only one concentration or points with standard deviations (SD) overlapping SD for control biofilm. However, a lot of isolates showed increase at one point or increase that did not put into our definition. So, the overall phenomenon might be broader than presented.

## Discussion

Clinical isolates in our study were obtained from the single hospital in Moscow, Russia. The general landscape of antibiotic resistance was similar to overall global picture with significant number of MDR *K. pneumoniae* in hospital setting. Whole-genome characterization of selected *K. pneumoniae* isolates showed diverse sequence types and variable combination of carbapenemase (KPC-3, NDM-1, OXA-48) and cephalosporinase-encoding (CTX-M-15) genes.

Considering lack of standardization in biofilm assays and any guidelines for *in vitro* screening assay we applied two technics – total biomass staining with crystal violet and metabolic activity measurement with resazurin. Both assays are common in fundamental biofilm research as useful approaches for screening purpose in contrast to labor-intensive CFU counting from biofilm. Crystal violet stains biofilm-embedded bacteria and matrix components, while resazurin is reduced to fluorescent resorufin via NADPH dehydrogenase-dependent electron transfer in metabolically active cells. In general, we obtained lower MBEC for biomass reduction than for metabolic suppression. In line with other studies MBEC for metabolic suppression were extremely high and unreachable *in vivo* in most cases. These data support knowledge of biofilm tolerance but may underestimate the relative efficiency of antibacterial therapy — a therapeutic goal that is hard to achieve, yet possible.

For the study of the effect on mature biofilms, antibiotics available in the Russian Federation were selected [31, 32, 33]. These antibiotics fall within the spectrum of natural activity against *K. pneumoniae*, are widely used for the treatment of community-acquired and hospital-acquired infections caused by this pathogen, and have a parenteral formulation [34, 35, 36]. Fluoroquinolones (ciprofloxacin, levofloxacin, moxifloxacin) – also drugs of choice (except for sepsis and septic shock) for infections caused by *Enterobacterales* resistant to third-generation cephalosporins (Watch group of the WHO AWaRe system); Carbapenems (meropenem, imipenem+cilastatin, and ertapenem) – as the group of choice for the treatment of severe infections caused by *Enterobacterales* resistant to third-generation cephalosporins (Watch group of the WHO AWaRe system). The drugs included in the study are as follows: Third-generation cephalosporins (ceftriaxone) – as the most commonly used group of antibiotics in Russian clinical practice: 5th place by Defined Daily Doses (DDDs) in the country and 1st place in the hospital sector, despite the increasing resistance to this group among *Enterobacterales* (Watch group of the WHO AWaRe system); Fourth-generation cephalosporins (cefepime) – as a drug potentially considered a carbapenem-sparing strategy (Watch group of the WHO AWaRe system); Aminoglycosides (amikacin and gentamicin) – which occupy a position as alternative or de-escalation therapy for urinary tract infections caused by *Enterobacterales* resistant to third-generation cephalosporins, or as a second agent in combination therapy for severe infections caused by carbapenem-resistant Gram-negative pathogens (Access group of the WHO AWaRe system) ; Ceftazidime/avibactam – as a first-line treatment for carbapenem-resistant *K. pneumoniae* infections not mediated by NDM production and/or non-enzymatic resistance mechanisms (Reserve group of the WHO AWaRe system); Colistimethate sodium – as a last-resort drug for the treatment of carbapenem-resistant *K. pneumoniae* infections (Reserve group of the WHO AWaRe system); Tigecycline – a reserve drug for use in combination therapy for infections caused by *K. pneumoniae* resistant to carbapenems and ceftazidime/avibactam (Reserve group of the WHO AWaRe system). With the exception of colistimethate, all selected antibiotics are included in the national EML. At the same time, the WHO EML does not include ertapenem, imipenem+cilastatin, cefepime, or tigecycline (the latter two were removed from the WHO EML in 2019). The following were not included among the tested drugs but are of no less interest in terms of their effect on mature *K. pneumoniae* biofilms: biapenem (due to experimental data indicating lower affinity for metallo-beta-lactamases), fosfomycin sodium salt, cefepime+sulbactam, and polymyxin B (all are included in the current national clinical guidelines and the national EML) [37]

All tested isolates were susceptible to colistin according to EUCAST and most responded to colistin sulfate in biofilm state. The pronounced activity of colistin against biofilms has been previously reported for *K. pneumonia* biofilms, as well as against *E. coli* biofilms and some non-enterobacteria species [17]. Thus, Wannigama showed that the difference between MIC and MBEC50 was the minimal for colistin (and the maximal for meropenem) in the case of both *P. aeruginosa* and *A. baumannii* biofilms [16]. Similar results were observed on isolates belonging to Liverpool Epidemic Strain (LES) of *P. aeruginosa* [38]. In the latter study polymixin B also has high activity against LES isolates, so polymixins as a class might have pronounce antibiofilm activity.

Tigecycline in general was able to reach MBEC50 for the vast majority of isolates. Tigecycline lacks activity against *P. aeruginosa*, but is the only agent with simultaneous activity against both ESBL-producing *Enterobacteriales* and MRSA. So, exploration of its antibiofilm activity especially interesting in the context of multispecies biofilms.

Notably, the pharmacokinetics (PK) of both tigecycline and colistin deviate from the common *C*_max_/MIC index used as a predictor of antibiotic efficacy. According to our data, if ignore plasma *C*_max_ they had the greatest chances of achieving both metabolic suppression and biomass eradication and might be considered as potent antibiofilm drugs. Other drugs rely on *C*_max_/MIC index, so their efficacy against biofilms should be interpreted with *C*_max_ in mind. So *C*_max_ as a reference point to cutoff therapeutically unreachable antibiotics concentrations were applied for the analysis.

Among studied cephalosporins cefepime and ceftriaxone were able to eradicate biomass in more than half cases, with the greatest potency of cefepime. Lee at al. have tested ceftazidime and cefepime in antibiotic lock technique (ALT) for the treatment of catheter-related infections. Under exposure to cefepime both two tested strains of *K. pneumonia* reduced CFU numbers to undetectable level by day 3-5 [39]. Tang et al. have reported the second lowest (after colistin) difference in MBEC/MIC for cefepime [28]. In other studies, cefepime alone or in combination with avibactame did not show significant antibiofilm activity against *Klebsiella* biofilm [40, 41]. Such contrast results must be additionally studied with different methodologies, exposure time and expanded number of isolates.

Fluoroquinolones and aminoglycosides had a less than 50% successful rate in reaching MBEC50 within concentration range below plasma *C*_max_. Ciprofloxacin has a tendency to be the less effective than moxifloxacin and levofloxacin.

Among the carbapenems, imipenem/cilastatin showed the greatest potency in biomass eradication. In a study of B. Veeraraghavan meropemen had MBEC from 2 to 128 mkg/ml for sensitive isolates and from 4 to 512 mkg/ml for resistant isolates, but sample size was only 8 isolates (4 S / 4 R) [21]. In study by M. Tang et al. imipenem was tested as solo compound (without cilastatin) and MBEC was measured as detected turbidity after regrowth (i.e., complete killing of every bacterium within biofilm). Imipenem «complete» MBEC was from 160 to 2560 mkg/ml [28]. In our study we did not observe good chances to reach metabolic suppression for carbapenemes while did not study extreme concentrations. According to our data carbapenemes have a very modest effect on mature biofilms: ertapenem and meropenem have reached biomass reduction in less than half sensitive isolates. Interestingly imipenem/cilastatin was more potent in biomass reduction in about 64% of isolates. The difference may be explained by differences in the binding of carbapenems to penicillin-binding proteins (PBPs). From one hand imipenem more strongly binds to PBP1a/PBP1b/PBP5 while meropenem binds stronger to PBP2/PBP3.

Elevated proportion of PBP5/6 (and PBP1a/1b) was shown for *Klebsilella* in comparison with *E. coli* [42]. In our previous proteomic study of clinical isolate of *P. aeruginosa* we have shown that PBPs is a part of extracellular biofilm matrix [9]. We hypothesize that *Klebsiella* biofilms might similarly overexpress PBPs (as carbapenem targets) or sequester them in the matrix as decoy molecules or as proteins with unexplored moonlight functions. On the other hand cilastatin is believed to inhibit human dehydropeptidase I, but in early publications it has been proved to be active against a zinc-dependent *β*-lactamase Cph1 from *Aeromonas hydrophila* as well [43]. For *Klebsiella* data on interaction of Cph1 homologs with cilastatin are missing. Further researches are needed to elucidate role of different PBPs in biofilms and possible cilastatin targets (*β*-lactamases?) in entrobacteriales. As to the secondary effects of carbapenems interesting data was reported by Chao et al. for carbapeneme resistant MDR *K. pneumonia*. According to the study carbapenemes enhance the secretion of outer membrane vesicles (OMVs) with altered protein composition such OMVs promote inflammation and decrease survival in mouse model of infection [44]. Among carbapenemes imipenem was the most potent agent in OMVs induction at least *in vitro*. Under exposure to carbapenemes composition of OMVs could have additional effect on biofilm biology and antibiotic susceptibility as OMVs are a part of biofilm matrix. Noteworthy antibiotic effect on OMVs not limited with carbapenemes, but also include other antibiotics and little studied under exposure to antibiotic supplements like cilastatin [45, 46].

MDR resistant *Klebsiella* in our study had a low level of resistance to colistin, tigecycline and ceftazidime/avibactam. Cefepime, ceftriaxone and imipenem/cilastatin despite activity against biofilms are usually have a resistant phenotype among MDR *K. pneumoniae*, so these antibiotics might be considered as a treatment option for sensitive phenotype and non-MDR strains.

Promotion of biofilm formation under stress condition is a well-known phenomenon. Exposure of bacterial suspension to antibiotics at subinhibitory concentration may cause switch to biofilm lifestyle as bacterial mechanism to defend external threat [47]. Logically exposure of biofilm to subMBEC of antibiotics may have secondary effects on biofilm.

Increase of CV-stained biomass of biofilms was reported for *P. aeruginosa* [38] Authors proposed hypothesis about extensive production of matrix components under subMBEC exposure. In another study biofilm morphology was affected by antibiotics and increase in cell volume and decrease in cell density precede bacterial death inside biofilm at least for *Vibrio cholerae* [48]. How it may influence biomass staining remains questionable.

Similarly, we found some tendance to increase of biomass staining for *K. pneumoniae*. Even more interesting was to found increase of metabolic activity at subMBEC of antibiotics. One possible explanation could be related to the biochemistry of the resazurin assay itself. According to microarray study antibiotic-induced response promote expression of NADPH dehydrogenase I gene in *E. coli* [49]. Overproduction of NADPH dehydrogenase may lead to inadequate interpretation of live bacteria content with resazurin as it is a key enzyme for electron transfer from NADH to resazurin. At the same time biochemical explanation does not exclude other «paradoxical effects». For example, Tsuji et al. have observed that higher polymyxin B exposures dramatically increased resistant subpopulations of *Acinetobacter baumannii* in the hollow-fiber infection model [50].

Resistant subpopulation might be actively metabolizing under subMBEC of antibiotics. The latter if translated to other bacteria have a direct consequence for antibiotic dose optimization and antimicrobial resistance arise.

Many studies rely on metrics directly or indirectly express total biomass content, including extracellular matrix, life and dead bacterial cells. While eradication of biomass does not express bacterial viability, but even if considered as «dead» biomass it still might be important target. At least *Vibrio cholerae* is able to recolonize dead biofilms[48]. So, biofilm eradication should be suggested as complex aim and includes both bacterial killing and biomass removal.

Bacterial subpopulations inside biofilm may differently respond to drugs, while CV staining and resazurin conversion express only cumulative data. With respect to well-known biofilm heterogeneity this question is out of scope of our study. As any technics have disadvantages, we consider to continue explore antibiofilm drugs with different assays.

Antibacterial therapy requires balance between toxicity, pharmacological interactions, immune status and not always matches with expectations based on *in vitro* studies.

Moreover, biofilms *in vivo* and *in vitro* have a different biology. It should be noted that antibiotics secondary effects on interconnected networks such as perturbation of normal microbiome and inflammation significantly affect therapy outcomes.

All of these put a limitation on the current study and must be evaluated in further researches. Nevertheless, all issues connected with antibiofilm activity should be looked as an argument to prioritize antibiotics for therapy of infections caused by biofilm forming *K. pneumoniae*.

## CONCLUSION

Generally, cefepime, ceftriaxone, imipenem/cilastatin, tigecycline and colistin sulfate had the ability to eradicate biofilm biomass *in vitro* in a concentration range below plasma (*C*_max_). Colistin is a reserved antibiotic. Cefepime, ceftriaxone and imipenem/cilastatin are classified in the ’Watch’ group of the WHO AWaRe classification. Ceftriaxone and colistin are in List of Essential Medicines. Among MDR *Klebsiella* in our study only colistin and tigecycline showed low level of resistance with simultaneous antibiofilm activity. Moreover, only colistin and tigecycline were able to reach MBEC in biomass eradication and in suppression of bacterial metabolic activity. Further research is needed for translation of *in vitro* data in clinical practice for prioritization of antibiotics in a case of biofilm and microbial aggregates. Current *in vitro* study might be considered along other studies in the field for knowledge-based prioritization of available antimicrobials in a case of limited number of drugs for therapy of infections caused by biofilm forming *K. pneumoniae*.

## Permission to use copyrighted material

Not applicable

## DATA AVAILABILITY STATEMENT

Clinical isolates are deposited in Gamaleya‘s collection of BSL1-2 pathogens. WGS data are available in GenBank: BioProject PRJNA561493, Accession Numbers are presented in supplementary Table 2.

## CLINICAL TRIALS

Not applicable

## ETHICS APPROVAL

This study was approved by the Institutional Review Board at N.F. Gamaleya NRCEM and Ethical committee at NMRC Center for Treatment and Rehabilitation (Approval Number: 38 from 17 March 2023)

## FUNDING

This research is supported by MoH of the Russian Federation (grant number 056-00121-26-01).

## CONFLICTS OF INTEREST

The authors declare no conflict of interest.

## Supplemental material

The following supporting information can be downloaded at article webpage: Figure S1: comparison of biofilm forming ability for different isolation source; Figure S2: MBEC for both R and S isolates; Table S1: S/R and MBEC table; Table S2: WGS data.

## References

[1] Høiby N, Bjarnsholt T, Moser C, Bassi GL, Coenye T, Donelli G, Hall-Stoodley L, Holá V, Imbert C, Kirketerp-Møller K, Lebeaux D, Oliver A, Ullmann AJ, Williams C. May 2015. ESCMID guideline for the diagnosis and treatment of biofilm infections 2014. Clinical Microbiology and Infection 21:S1–S25. doi:10.1016/j.cmi.2014.10.024.

[2] Lichtenberg M, Coenye T, Parsek MR, Bjarnsholt T, Jakobsen TH. Sep 2023. What’s in a name? Characteristics of clinical biofilms. FEMS Microbiology Reviews 47 (5):fuad050. doi:10.1093/femsre/fuad050.

[3] Liu Y, Pan C, Ye L, Si Y, Bi C, Hua X, Yu Y, Zhu L, Wang H. Jun 2020. Nonclassical Biofilms Induced by DNA Breaks in Klebsiella pneumoniae. mSphere 5 (3):e00336–20. doi:10.1128/mSphere.00336-20.

[4] Kolpen M, Kragh KN, Enciso JB, Faurholt-Jepsen D, Lindegaard B, Egelund GB, Jensen AV, Ravn P, Mathiesen IHM, Gheorge AG, Hertz FB, Qvist T, Whiteley M, Jensen PØ, Bjarnsholt T. Oct 2022. Bacterial biofilms predominate in both acute and chronic human lung infections. Thorax 77 (10):1015–1022. doi: 10.1136/thoraxjnl-2021-217576.

[5] Ciofu O, Moser C, Jensen PØ, Høiby N. Oct 2022. Tolerance and resistance of microbial biofilms. Nature Reviews Microbiology 20 (10):621–635. doi: 10.1038/s41579-022-00682-4.

[6] Hall CW, Mah TF. May 2017. Molecular mechanisms of biofilm-based antibiotic resistance and tolerance in pathogenic bacteria. FEMS Microbiology Reviews 41 (3):276–301. doi:10.1093/femsre/fux010.

[7] Chiang WC, Nilsson M, Jensen PØ, Høiby N, Nielsen TE, Givskov M, Tolker Nielsen T. May 2013. Extracellular DNA Shields against Aminoglycosides in *Pseudomonas aeruginosa* Biofilms. Antimicrobial Agents and Chemotherapy 57 (5):2352–2361. doi:10.1128/AAC.00001-13.

[8] Park AJ, Surette MD, Khursigara CM. Sep 2014. Antimicrobial targets localize to the extracellular vesicle-associated proteome of *Pseudomonas aeruginosa* grown in a biofilm. Frontiers in Microbiology 5. doi:10.3389/fmicb.2014.00464.

[9] Egorova DA, Solovyev AI, Polyakov NB, Danilova KV, Scherbakova AA, Kravtsov IN, Dmitrieva MA, Rykova VS, Tutykhina IL, Romanova YM, Gintsburg AL. Sep 2022. Biofilm matrix proteome of clinical strain of *P. aeruginosa* isolated from bronchoalveolar lavage of patient in intensive care unit. Microbial Pathogenesis 170:105714. doi:10.1016/j.micpath.2022.105714.

[10] Bano S, Hassan N, Rafiq M, Hassan F, Rehman M, Iqbal N, Ali H, Hasan F, Kang YQ. Oct 2023. Biofilms as Battlefield Armor for Bacteria against Antibiotics: Challenges and Combating Strategies. Microorganisms 11 (10):2595. doi: 10.3390/microorganisms11102595.

[11] Prax M, Bertram R. Oct 2014. Metabolic aspects of bacterial persisters. Frontiers in Cellular and Infection Microbiology 4. doi:10.3389/fcimb.2014.00148.

[12] Tahrioui A, Duchesne R, Bouffartigues E, Rodrigues S, Maillot O, Tortuel D, Hardouin J, Taupin L, Groleau MC, Dufour A, Déziel E, Brenner-Weiss G, Feuilloley M, Orange N, Lesouhaitier O, Cornelis P, Chevalier S. May 2019. Extracellular DNA release, quorum sensing, and PrrF1/F2 small RNAs are key players in *Pseudomonas aeruginosa* tobramycin-enhanced biofilm formation. npj Biofilms and Microbiomes 5 (1):15. doi:10.1038/s41522-019-0088-3.

[13] Nesse LL, Simm R. 2018. Biofilm: A Hotspot for Emerging Bacterial Genotypes. Advances in Applied Microbiology 103:223–246. doi:10.1016/bs.aambs.2018.01.003.

[14] Høiby N, Moser C, Oliver A, Williams C, Ramage G, Borghi E, Azeredo J, Macia MD. 2023. To update or not to update the ESCMID guidelines for the diagnosis and treatment of biofilm infections – That is the question! The opinion of the ESGB board. Biofilm 6:100135. 10.1016/j.bioflm.2023.100135.

[15] Okae Y, Nishitani K, Sakamoto A, Kawai T, Tomizawa T, Saito M, Kuroda Y, Matsuda S. Jul 2022. Estimation of Minimum Biofilm Eradication Concentration (MBEC) on In Vivo Biofilm on Orthopedic Implants in a Rodent Femoral Infection Model. Frontiers in Cellular and Infection Microbiology 12:896978. doi: 10.3389/fcimb.2022.896978.

[16] Wannigama DL, Hurst C, Hongsing P, Pearson L, Saethang T, Chantaravisoot N, Singkham-in U, Luk-in S, Storer RJ, Chatsuwan T. Dec 2020. A rapid and simple method for routine determination of antibiotic sensitivity to biofilm populations of *Pseudomonas aeruginosa*. Annals of Clinical Microbiology and Antimicrobials 19 (1):8. doi: 10.1186/s12941-020-00350-6.

[17] Klinger-Strobel M, Stein C, Forstner C, Makarewicz O, Pletz MW. Apr 2017. Effects of colistin on biofilm matrices of *Escherichia coli* and *Staphylococcus aureus*. International Journal of Antimicrobial Agents 49 (4):472–479. doi:10.1016/j.ijantimicag.2017.01.005.

[18] Ramanan L, Ragupathi NKD, Sethuvel DPM, Balaji V. Mar 2022. MBEC Determination Guides to Choose the Optimal Drug And Dose in Biofilm Forming Hypervirulent *Klebsiella pneumoniae*: Value Addition to Antimicrobial Stewardship. International Journal of Infectious Diseases 116:S10–S11. doi:10.1016/j.ijid.2021.12.025.

[19] Singla S, Harjai K, Chhibber S. Feb 2013. Susceptibility of different phases of biofilm of *Klebsiella pneumoniae* to three different antibiotics. The Journal of Antibiotics 66 (2):61–66. doi:10.1038/ja.2012.101.

[20] Anderl JN, Zahller J, Roe F, Stewart PS. Apr 2003. Role of Nutrient Limitation and Stationary-Phase Existence in *Klebsiella pneumoniae* Biofilm Resistance to Ampicillin and Ciprofloxacin. Antimicrobial Agents and Chemotherapy 47 (4):1251–1256. doi: 10.1128/AAC.47.4.1251-1256.2003.

[21] Devanga Ragupathi NK, Muthuirulandi Sethuvel DP, Triplicane Dwarakanathan H, Murugan D, Umashankar Y, Monk PN, Karunakaran E, Veeraraghavan B. Dec 2020. The Influence of Biofilms on Carbapenem Susceptibility and Patient Outcome in Device Associated K. pneumoniae Infections: Insights Into Phenotype vs Genome-Wide Analysis and Correlation. Frontiers in Microbiology 11:591679. doi: 10.3389/fmicb.2020.591679.

[22] Høiby N, Moser C, Oliver A, Williams C, Ramage G, Borghi E, Azeredo J, Macia MD. Dec 2023. To update or not to update the ESCMID guidelines for the diagnosis and treatment of biofilm infections – That is the question! The opinion of the ESGB board. Biofilm 6:100135. doi:10.1016/j.bioflm.2023.100135.

[23] Razdan K, Garcia-Lara J, Sinha VR, Singh KK. Aug 2022. Pharmaceutical strategies for the treatment of bacterial biofilms in chronic wounds. Drug Discovery Today 27 (8):2137–2150. doi:10.1016/j.drudis.2022.04.020.

[24] Ferreira M, Pinto M, Aires-da Silva F, Bettencourt A, Gaspar MM, Aguiar SI. Oct 2024. Rifabutin: a repurposed antibiotic with high potential against planktonic and biofilm staphylococcal clinical isolates. Frontiers in Microbiology 15:1475124. doi: 10.3389/fmicb.2024.1475124.

[25] De Oliveira DMP, Forde BM, Kidd TJ, Harris PNA, Schembri MA, Beatson SA, Paterson DL, Walker MJ. Jun 2020. Antimicrobial Resistance in ESKAPE Pathogens. Clinical Microbiology Reviews 33 (3):e00181–19. doi:10.1128/CMR.00181-19.

[26] Naparstek L, Carmeli Y, Navon-Venezia S, Banin E. Apr 2014. Biofilm formation and susceptibility to gentamicin and colistin of extremely drug-resistant KPC-producing *Klebsiella pneumoniae*. Journal of Antimicrobial Chemotherapy 69 (4):1027–1034. doi:10.1093/jac/dkt487.

[27] Shein AMS, Wannigama DL, Higgins PG, Hurst C, Abe S, Hongsing P, Chantaravisoot N, Saethang T, Luk-in S, Liao T, Nilgate S, Rirerm U, Kueakulpattana N, Laowansiri M, Srisakul S, Muhummudaree N, Techawiwattanaboon T, Gan L, Xu C, Kupwiwat R, Phattharapornjaroen P, Rojanathanes R, Leelahavanichkul A, Chatsuwan T. Nov 2021. Novel colistin-EDTA combination for successful eradication of colistin-resistant *Klebsiella pneumoniae* catheter-related biofilm infections. Scientific Reports 11 (1):21676. doi: 10.1038/s41598-021-01052-5.

[28] Tang M, Wei X, Wan X, Ding Z, Ding Y, Liu J. Oct 2020. The role and relationship with efflux pump of biofilm formation in *Klebsiella pneumoniae*. Microbial Pathogenesis 147:104244. doi:10.1016/j.micpath.2020.104244.

[29] Magréault S, Jauréguy F, Carbonnelle E, Zahar JR. 2022. When and How to Use MIC in Clinical Practice? Antibiotics 11 (12). doi:10.3390/antibiotics11121748.

[30] Tsuji BT, Pogue JM, Zavascki AP, Paul M, Daikos GL, Forrest A, Giacobbe DR, Viscoli C, Giamarellou H, Karaiskos I, Kaye D, Mouton JW, Tam VH, Thamlikitkul V, Wunderink RG, Li J, Nation RL, Kaye KS. Jan 2019. International Consensus Guidelines for the Optimal Use of the Polymyxins: Endorsed by the American College of Clinical Pharmacy (ACCP), European Society of Clinical Microbiology and Infectious Diseases (ESCMID), Infectious Diseases Society of America (IDSA), International Society for Anti-infective Pharmacology (ISAP), Society of Critical Care Medicine (SCCM), and Society of Infectious Diseases Pharmacists (SIDP). Pharmacotherapy: The Journal of Human Pharmacology and Drug Therapy 39 (1):10–39. doi: 10.1002/phar.2209.

[31] Arepyeva MA, Kuzmenkov AY, Starostenkov AA, Kolbin AS, Balykina YE, Yulia M, Kurylev AA, Kozlov RS. Jan 2026. An integrated monitoring system for antimicrobial consumption and resistance forecasting using dynamic models. Clinical Microbiology and Antimicrobial Chemotherapy 27 (3):330. doi: 10.36488/cmac.2025.3.330-341.

[32] Arepyeva MA, Kuzmenkov AY, Starostenkov AA, Kolbin AS, Balykina YE, Gomon YM, Kurylev AA, Kozlov RS, Sidorenko SV. Feb 2026. Predictive modelling of the dynamics of antimicrobial resistance: creation of a bank of renewable models based on machine learning. Frontiers in Pharmacology 17:1715346. doi: 10.3389/fphar.2026.1715346.

[33] Gomon YM, Kolbin AS, Arepyeva MA, Kalyapin AA, Balykina YE, Kurylev AA, Kuzmenkov AY, Kozlov RS. 2023. Antimicrobial drug consumption in the Russian Federation (2008–2022): pharmacoepidemiological study. Clinical Microbiology and Antimicrobial Chemotherapy 25 (4):395–400. doi:10.36488/cmac.2023.4.395-400.

[34] Paul M, Carrara E, Retamar P, Tängdén T, Bitterman R, Bonomo RA, De Waele J, Daikos GL, Akova M, Harbarth S, Pulcini C, Garnacho-Montero J, Seme K, Tumbarello M, Lindemann PC, Gandra S, Yu Y, Bassetti M, Mouton JW, Tacconelli E, Rodríguez-Baño J. Apr 2022. European Society of Clinical Microbiology and Infectious Diseases (ESCMID) guidelines for the treatment of infections caused by multidrug-resistant Gram-negative bacilli (endorsed by European society of intensive care medicine). Clinical Microbiology and Infection 28 (4):521–547. doi: 10.1016/j.cmi.2021.11.025.

[35] Tamma PD, Heil EL, Justo JA, Mathers AJ, Satlin MJ, Bonomo RA. Aug 2024. Infectious Diseases Society of America 2024 Guidance on the Treatment of Antimicrobial-Resistant Gram-Negative Infections. Clinical Infectious Diseases p ciae403. doi:10.1093/cid/ciae403.

[36] Beloborodov VB, Goloshchapov OV, Gusarov VG, Dekhnich AV, Zamyatin MN, Zolotukhin KN, Zubareva NA, Zyryanov SK, Kamyshova DA, Klimko NN, Kozlov RS, Kulabukhov VV, Matinyan NV, Petrushin MA, Polushin YS, Popov DA, Pyregov AV, Rudnov VA, Sidorenko SV, Sokolov DV, Sychev IN, Shlyk IV, Eydelstein MV, Yakovlev SV. Apr 2025. Diagnosis and antimicrobial therapy of infections caused by polyresistant microorganisms (updated 2024). Messenger of ANESTHESIOLOGY AND RESUSCITATION 22 (2):149–189. doi: 10.24884/2078-5658-2025-22-2-149-189.

[37] Tapalskiy DV, Karpova EV, Simonchik MV, Pyzh AE, Golikova MV. Jan 2026. Comparative activity of meropenem and biapenem and their combinations with colistin against Gram-negative microorganisms-producers of carbapenemases from various groups. Clinical Microbiology and Antimicrobial Chemotherapy 27 (3):317. doi: 10.36488/cmac.2025.3.317-329.

[38] Goodyear MC, Garnier NE, Levesque RC, Khursigara CM. Jun 2022. Liverpool Epidemic Strain Isolates of *Pseudomonas aeruginosa* Display High Levels of Antimicrobial Resistance during Both Planktonic and Biofilm Growth. Microbiology Spectrum 10 (3):e01024–22. doi:10.1128/spectrum.01024-22.

[39] Lee MY, Ko KS, Song JH, Peck KR. Jul 2007. In vitro effectiveness of the antibiotic lock technique (ALT) for the treatment of catheter-related infections by Pseudomonas aeruginosa and Klebsiella pneumoniae. Journal of Antimicrobial Chemotherapy 60 (4):782–787. doi:10.1093/jac/dkm295.

[40] Shipitsyna IV, Osipova EV. Dec 2022. Resistance of *Klebsiella* pneumoniae isolated from patients with chronic osteomyelitis to antibacterial drugs. Kazan medical journal 103 (6):962–967. doi:10.17816/KMJ88660.

[41] Tian M, Yan B, Jiang R, Liu C, Li Y, Xu B, Guo S, Li X. Oct 2024. Activity of polymyxin B combined with cefepime-avibactam against the biofilms of polymyxin B-resistant *Pseudomonas* aeruginosa and *Klebsiella pneumoniae* in in vitro and in vivo models. BMC Microbiology 24 (1):409. doi:10.1186/s12866-024-03571-3.

[42] Sutaria DS, Moya B, Green KB, Kim TH, Tao X, Jiao Y, Louie A, Drusano GL, Bulitta JB. Jun 2018. First Penicillin-Binding Protein Occupancy Patterns of *β*-Lactams and *β*-Lactamase Inhibitors in *Klebsiella pneumoniae*. Antimicrobial Agents and Chemotherapy 62 (6):e00282–18. doi:10.1128/AAC.00282-18.

[43] Keynan S, Hooper NM, Felici A, Amicosante G, Turner AJ. Jul 1995. The renal membrane dipeptidase (dehydropeptidase I) inhibitor, cilastatin, inhibits the bacterial metallo-beta-lactamase enzyme CphA. Antimicrobial Agents and Chemotherapy 39 (7):1629–1631. doi:10.1128/AAC.39.7.1629.

[44] Ye C, Li W, Yang Y, Liu Q, Li S, Zheng P, Zheng X, Zhang Y, He J, Chen Y, Hua L, Yang Z, Li D, Ren Z, Yang Y, Qi J, Huang W, Ma Y. Sep 2021. Inappropriate use of antibiotics exacerbates inflammation through OMV-induced pyroptosis in MDR *Klebsiella pneumoniae* infection. Cell Reports 36 (12):109750. doi: 10.1016/j.celrep.2021.109750.

[45] Hussein M, Jasim R, Gocol H, Baker M, Thombare VJ, Ziogas J, Purohit A, Rao GG, Li J, Velkov T. Feb 2023. Comparative Proteomics of Outer Membrane Vesicles from Polymyxin-Susceptible and Extremely Drug-Resistant *Klebsiella pneumoniae*. mSphere 8 (1):e00537–22. doi:10.1128/msphere.00537-22.

[46] Lucena ACR, Ferrarini MG, De Oliveira WK, Marcon BH, Morello LG, Alves LR, Faoro H. May 2023. Modulation of *Klebsiella pneumoniae* Outer Membrane Vesicle Protein Cargo under Antibiotic Treatment. Biomedicines 11 (6):1515. doi: 10.3390/biomedicines11061515.

[47] Ranieri MR, Whitchurch CB, Burrows LL. Oct 2018. Mechanisms of biofilm stimulation by subinhibitory concentrations of antimicrobials. Current Opinion in Microbiology 45:164–169. doi:10.1016/j.mib.2018.07.006.

[48] Díaz-Pascual F, Hartmann R, Lempp M, Vidakovic L, Song B, Jeckel H, Thormann KM, Yildiz FH, Dunkel J, Link H, Nadell CD, Drescher K. Oct 2019. Breakdown of *Vibrio cholerae* biofilm architecture induced by antibiotics disrupts community barrier function. Nature Microbiology 4 (12):2136–2145. doi:10.1038/s41564-019-0579-2.

[49] Kohanski MA, Dwyer DJ, Hayete B, Lawrence CA, Collins JJ. Sep 2007. A Common Mechanism of Cellular Death Induced by Bactericidal Antibiotics. Cell 130 (5):797–810. doi:10.1016/j.cell.2007.06.049.

[50] Tsuji BT, Landersdorfer CB, Lenhard JR, Cheah SE, Thamlikitkul V, Rao GG, Holden PN, Forrest A, Bulitta JB, Nation RL, Li J. Jul 2016. Paradoxical Effect of Polymyxin B: High Drug Exposure Amplifies Resistance in *Acinetobacter baumannii*. Antimicrobial Agents and Chemotherapy 60 (7):3913–3920. doi:10.1128/AAC.02831-15.

